# The Prophylactic and Therapeutic Efficacy of the Broadly Active Antiviral Ribonucleoside *N*^*4*^-Hydroxycytidine (EIDD-1931) in a Mouse Model of Lethal Ebola Virus Infection

**DOI:** 10.1101/2022.10.18.512767

**Authors:** Gregory R. Bluemling, Shuli Mao, Michael G. Natchus, Wendy Painter, Sabue Mulangu, Mark Lockwood, Abel De La Rosa, Trevor Brasel, Jason E. Comer, Alexander N. Freiberg, Alexander A. Kolykhalov, George R. Painter

## Abstract

The unprecedented magnitude of the 2013-2016 Ebola virus (EBOV) epidemic in West Africa resulted in over 11,000 deaths and spurred an international public health emergency. A second outbreak in 2018-2020 in DRC resulted in an additional >3400 cases and nearly 2300 deaths (WHO Disease Outbreak News: Update 26 June, 2020). These large outbreaks across geographically diverse regions highlight the need for the development of effective oral therapeutic agents that can be easily distributed for self-administration to populations with active disease or at risk of infection. Herein, we report the in vivo efficacy of *N*^*4*^-hydroxycytidine (EIDD-1931), a broadly active ribonucleoside analog and the active metabolite of the prodrug EIDD-2801 (molnupiravir), in murine models of lethal EBOV infection. Twice daily oral dosing with EIDD-1931 at 200 mg/kg for 7 days, initiated either with a prophylactic dose 2 hours before infection, or as therapeutic treatment starting 6 hours post-infection, resulted in 92-100% survival of mice challenged with lethal doses of EBOV, reduced clinical signs of Ebola virus disease (EVD), reduced serum virus titers, and facilitated weight loss recovery. These results support further investigation of molnupiravir as a potential therapeutic or prophylactic treatment for EVD.

## 1. INTRODUCTION

At least 30 outbreaks of Ebola virus disease (EVD) have occurred since the late 1970s with the vast majority occurring in isolated rural villages (Keita et al. 2021). However, the 2013-2016 and 2018-2020 outbreaks demonstrated that EBOV can be spread easily in densely populated urban centers. Moreover, owing to rapid international travel, the disease could be further spread to populations that had not been at risk of infection with this previously geographically limited virus. The EBOV outbreak in Guinea in 2021 added a new dimension to how disease can be transmitted. Up until this outbreak it had been assumed that all new outbreaks were the result of independent zoonotic transmissions from intermediate or amplifying animal reservoirs and then continued via human-to-human spread by direct contact with contaminated bodily fluids and tissues (Rewar & Mirdha 2014). However, genomic analysis of virus isolated from patients in the 2021 outbreak strongly suggests that this virus was related to the one causing the 2013-2016 outbreak, and that cases in the later outbreak were likely the result of persistent infection in a former survivor. The persistent infection led to subsequent transmission of the virus (Keita et al. 2021).

EVD is severe and often fatal. A large percentage of patients fall ill within 2 to 20 days after infection with a 50 to 90% death rate observed from the most pathogenic EBOV species. During the early stage of EVD, patients usually present with non-specific flu-like symptoms that rapidly progress to gastrointestinal symptoms such as vomiting and diarrhea (Feldmann & Geisbert 2011). During the late stage of disease, there is an increase in vascular permeability and a systemic inflammatory response with massive tissue injury. At this stage, hemorrhagic manifestations such as petechiae, ecchymosis, and mucosal hemorrhages can be seen (Kortepeter et al. 2011). The main causes of death are multiple organ failure and shock (Bwaka et al. 1999). In survivors, EBOV can persist in various immunologically protected body compartments and fluids such as semen, chambers of the eye, and the central nervous system for months after infection. The persisting virus can lead to EVD-related sequelae, transmission, and virus reactivation (Vetter et al. 2016a, Vetter et al. 2016b). Sexual transmission from convalescent survivors likely occurs and is a suspected cause of re-emergence (Den Boon et al. 2019). Given the severity of disease, high fatality rate, and extreme threat to public health that Ebola presents, the National Institute of Allergy and Infectious Diseases (NIAID) and the Centers for Disease Control (CDC) have classified it as a Category A pathogen.

Two Ebola vaccines have been approved (EMA 2019, FDA 2019). However, the strategy for how best to use them across the endemic range of the disease is still being debated given the infrequency of outbreaks, as well as socio-economic and political environments (Bausch 2021). The use of vaccines in non-endemic regions to protect against spread by infected travelers or for protection against a bioterrorist attack also seems unlikely.

Two monoclonal antibody (mAb)-based therapeutics have been approved for the treatment of EBOV infection: Ebanga™ (ansuvimab-zykl) and Inmazeb™ (a mixture of maftivimab, odesivimab, and atoltivimab). Though effective, the intravenous administration of monoclonal therapies requires specialized medical infrastructure which are often not available in rural areas. Additionally, limited penetration of antibodies into immune sanctuaries where EBOV continues to replicate has been reported, and this can result in relapses and persistent infection leading to transmission (Keita et al, 2021). Currently, there are no FDA approved oral therapeutic agents for the treatment or prevention of EBOV infection in humans. Given the issues associated with vaccination and the medical infrastructure required for the intravenous administration of monoclonal therapies, it is clear that oral therapeutics, that can be self-administered, are needed for treatment and post-exposure prophylaxis. They are important not only in rural settings, where social and logistic issues limit the effectiveness of vaccines and monoclonal therapies, but also in densely populated areas where large numbers of people can be exposed and infected quickly.

They are also needed for non-endemic areas where the disease may be introduced by an infected international traveler or by a bioterrorist attack. Here we report the prophylactic and therapeutic efficacy of orally administered N4-hydroxycytidine (NHC), also identified as EIDD-1931 (**Fig. 1**), in a murine model of lethal EBOV in which infection is initiated by intraperitoneal injection of a mouse adapted strain of EBOV. This model provides an effective means of establishing in vivo efficacy; it is characterized by defined clinical signs, uniform lethality, and a narrow window for time to death (Haddock et al. 2018). Additionally, the cellular hallmarks of disease in mice are very similar to the pathogenesis observed in the nonhuman primate (NHP) model of disease (Taylor 2007). The results of the in vivo efficacy studies reported here strongly support the continued development of EIDD-1931 for the prophylaxis and treatment of EVD.

**Figure 1:**
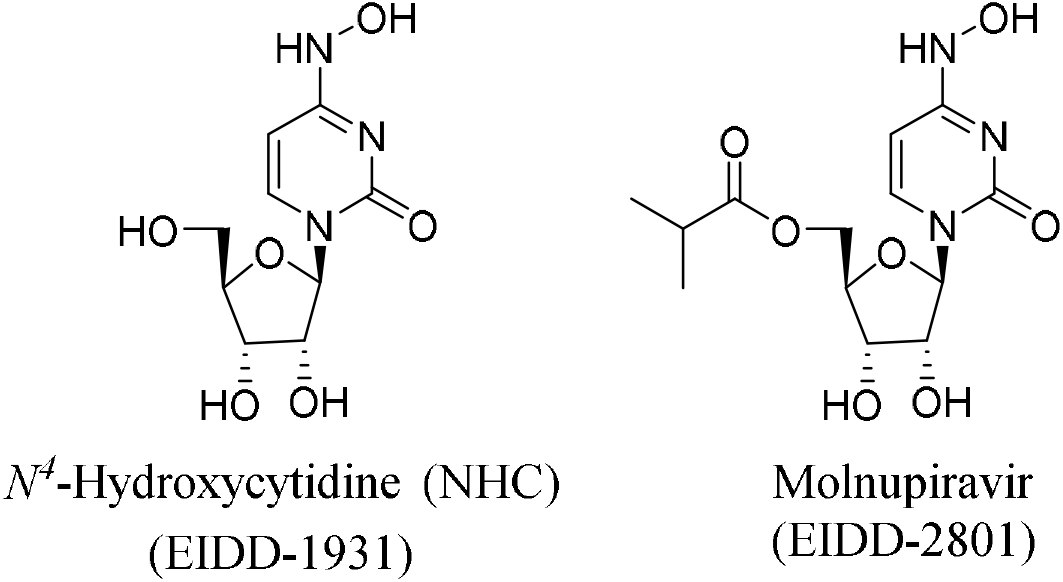
EIDD-1931, N^4^-hydroxycytidine, after phosphorylation by host kinases acts as a competitive, alternative substrate for the virally encoded RNA-dependent RNA-polymerase (Yoon et al. 2018, Painter et al. 2019, Gordon et al. 2021). Incorporation of EIDD-1931-5’-monophosphate into nascent chain RNA has been shown to cause acceleration of viral decay (“error catastrophe”) and loss of virulence (Urakova et al. 2018, Yoon et al. 2018, Agostini et al. 2019, Sheahan et al. 2020). Molnupiravir (EIDD-2801), which has been authorized for emergency use in multiple countries for the treatment of mild to moderate COVID-19, is the 5’-isobutyrate ester prodrug of EIDD-1931.

## 2. Materials and Methods

### Compounds and Viruses

EIDD-1931, greater than 97% pure, was prepared at the Emory Institute for Drug Development as previously described (Painter et al. 2019). A mouse-adapted Ebola virus (maEBOV; *M*.*musculus*/COD/1976/Mayinga-CDC-808012), originally derived from the 1976 Ebola virus Zaire isolate Yambuku-Maynga (Bray et al. 1998), was propagated at the University of Texas Medical Branch (UTMB) and used in all in vivo experiments. All work with infectious virus was conducted under Biological Safety Level 4 (BSL-4) conditions in the Robert E. Shope Laboratory and Galveston National Laboratory BSL-4 laboratories at the UTMB.

### In vivo Efficacy testing

All animal experiments were approved by the Institutional Animal Care and Use Committee of The University of Texas Medical Branch and performed following the guidelines of the Association for Assessment and Accreditation of Laboratory Animal Care International (AAALAC) by certified staff in an AAALAC-approved facility.

BALB/c mice (equal sex, 8-9 weeks old at the time of infection) were obtained from Envigo Laboratories and housed under specific pathogen-free conditions. Food, water, and bedding were routinely supplied, changed, and monitored. Mice were assigned to study groups as shown in **Table 1**.

**TABLE 1.** Experimental Design for Treating Ebola-Challenged Mice with EIDD-1931

| Study <sup>a</sup> | Group | n <sup>b</sup> | Treatment | Dose Level | Dose Volume | Route <sup>c</sup> | Dosing Initiation and Frequency <sup>d</sup> |
| --- | --- | --- | --- | --- | --- | --- | --- |
| 1 | 1 | 12 | Vehicle <sup>e</sup> | - | 0.1 mL | PO | On Day 0, dosing occurred 2h ± 15 min prior to virus challenge, 4h ± 30 min post-challenge, then QD (24h ± 1h, throughout) for 6 additional days (n=8 total doses) |
|  | 2 | 12 | EIDD-1931 | 100 mg/kg | 0.1 mL | PO |  |
|  | 3 | 12 | EIDD-1931 | 200 mg/kg | 0.2 mL | PO |  |
|  | 4 | 12 | EIDD-1931 | 200 mg/kg | 0.2 mL | PO | On Day 0, dosing occurred 2h ± 15 min prior to virus challenge, 4h ± 30 min post-challenge, 12h ± 1h post-challenge, then BID (12h ± 1h, relevant to the challenge timepoint) for 6 additional days (n=15 total doses) |
| 2 | 1 | 12 | Vehicle <sup>e</sup> | - | 0.1 mL | PO | BID (12h ± 1h apart) beginning 6h ± 30min post-challenge on Day 0 for 7 days (14 total doses) |
|  | 2 | 12 | EIDD-1931 | 200 mg/kg | 0.2 mL | PO |  |
|  | 3 | 12 | EIDD-1931 | 200 mg/kg | 0.2 mL | PO | BID (12h ± 1h apart) beginning 24h ± 1h post-challenge on Day 0 for 7 days (14 total doses) |

<sup>a</sup> prophylactic treatment (Study 1) and therapeutic treatment (Study 2)
<sup>b</sup> n, number of animals; equal sex
<sup>c</sup> PO, *per os* via oral gavage
<sup>d</sup> QD, daily; BID, twice daily
<sup>e</sup> mice treated with vehicle only (0.24M sodium citrate, pH 3)

Mice were challenged with a target dose of 100 PFU maEBOV via intraperitoneal injection. Virus challenge dose was verified using a standard plaque assay as previously described (Shurtleff et al. 2016). Animals were dosed with vehicle or EIDD-1931 per os (0.1 to 0.2 mL as indicated, **Table 1**) following sedation with isoflurane. In addition to the listed groups, two sham-infected control groups were used in each study that received either vehicle or the highest dose of EIDD-1931 (200 mg/kg). Animals were monitored daily for development of clinical signs and changes in body weight. Clinical scoring and health assessments were performed and documented at each observation. The following scoring chart was used: 1, healthy; 2, displaying mild signs of lethargy, some fur ruffling, no hunched posture; 3, fur ruffling, hunched posture, mild signs of lethargy; 4, ruffled, hunched posture, increased lethargy and limited mobility; 5 (immediate euthanasia), moribund, ruffled, hunched posture with reduced or minimal mobility consistent with inability to reach food or water OR ≥20% weight loss. In cases where several observations occurred on the same day, clinical score was calculated by averaging the observations. Body weights were monitored daily through day 10 post-challenge then every third day until the end of the study.

### Terminal Serum Viral Load Measurement

When possible, blood was collected into serum separator tubes at terminal euthanasia for viral load measurement via quantitative real-time PCR (qRT-PCR) and plaque assay. Collected sera for qRT-PCR analysis was added to TRIzol® LS Reagent (Thermo Fisher, Waltham, MA, USA) (1:5 ratio), mixed, then stored frozen at −80°C. For analysis, samples were processed for RNA extraction and purification using the Zymo Direct-zol™ RNA Mini Prep kit (Zymo Research, Irvine, CA, USA) per manufacturer instructions. RNA samples were reverse transcribed and amplified using TaqMan™ chemistry and the CFX96 Touch™ Real-Time PCR Detection System (Bio-Rad, Hercules, CA, USA). Primers and probe targeted the glycoprotein gene from EBOV (GenBank accession no. **AF086833**). For quantification purposes, an HPLC-purified synthetic EBOV RNA containing the conserved EBOV glycoprotein sequence was used.

Collected sera for plaque assay analysis was stored frozen at −80°C. For analysis, serum samples were r serially diluted in filter-sterilized dilution medium (MEM/1% heat-inactivated fetal bovine serum/1% Penicillin-Streptomycin) and were analyzed according to standard plaque assay on Vero E6 cell monolayers as previously described (Shurtleff et al. 2016).

## 3. RESULTS

Given the high exposures achievable in mice plasma and organs after oral dosing (Painter et al. 2019), the in vitro efficacy of EIDD-1931 (3-3.8 µM EC_50_; Reynard et al. 2015) was deemed sufficient to justify testing in a mouse model of Ebola infection. EIDD-1931 was first tested as a prophylaxis for lethal EBOV infection in BALB/c mice in which dosing was initiated 2 hours prior to infection. This study was followed by a therapeutic treatment study in which dosing was initiated 6 or 24 hours post infection. Mice were challenged by an intraperitoneal injection of approximately 100 plaque forming units (PFU) of mouse-adapted EBOV virus (maEBOV), an exposure that has previously been shown to be 100% lethal in this model (Bray et al. 1998). In the prophylaxis study the challenge dose was confirmed to be between 80 and 116 PFU; in the therapeutic study the challenge dose was verified to be higher, between 244 and 273 PFU. Each study had an infected control group that received vehicle only as well as two sham-infected controls that received either vehicle or the highest dose of EIDD-1931 used in the study. Consistent with previous studies (Painter et al. 2019), none of the EIDD-1931 treated sham-infected mice exhibited any signs of toxicity as evidenced by lack of weight loss or increase in clinical scores (data not shown).

### 3.1 Prophylactic Dosing with EIDD-1931

On study Day 0, the same prophylactic dose of 200 mg/kg of EIDD-1931 was administered 2 hours prior to virus challenge to three groups of mice (Groups 2-4; **Table 1**, Study 1). Mice in groups 2 and 3 were then administered 4 hours post virus challenge either a 100 or 200 mg/kg dose of EIDD-1931, respectively, then once daily (QD) starting 24 hours post-challenge with the same doses for 6 additional days (8 total treatment doses). Mice in group 4 were administered a 200 mg/kg dose of EIDD-1931 4 and 12 hours post virus challenge and then treated BID (every 12 hours) with the same dose for six additional days (15 total treatment doses). A vehicle treatment group (group 1) of maEBOV-challenged mice was dosed QD for 6 days. The Kaplan-Meier survival curves for the prophylactic study are shown in **Fig. 2A**. All mice challenged with virus that received daily administration of vehicle succumbed on Study Days 3-7, the expected time to death following administration of 100 PFU (approx. 3000 LD_50_) EBOV (Bray et al. 1998). Mice challenged with virus that received QD administration of EIDD-1931 at 100 mg/kg or 200 mg/kg demonstrated 50% and 58.3% survival rates, respectively, p<0.0001 for both. All mice that received 200 mg/kg EIDD-1931 BID survived to the end of the 24-day study period.

**FIG 2.**
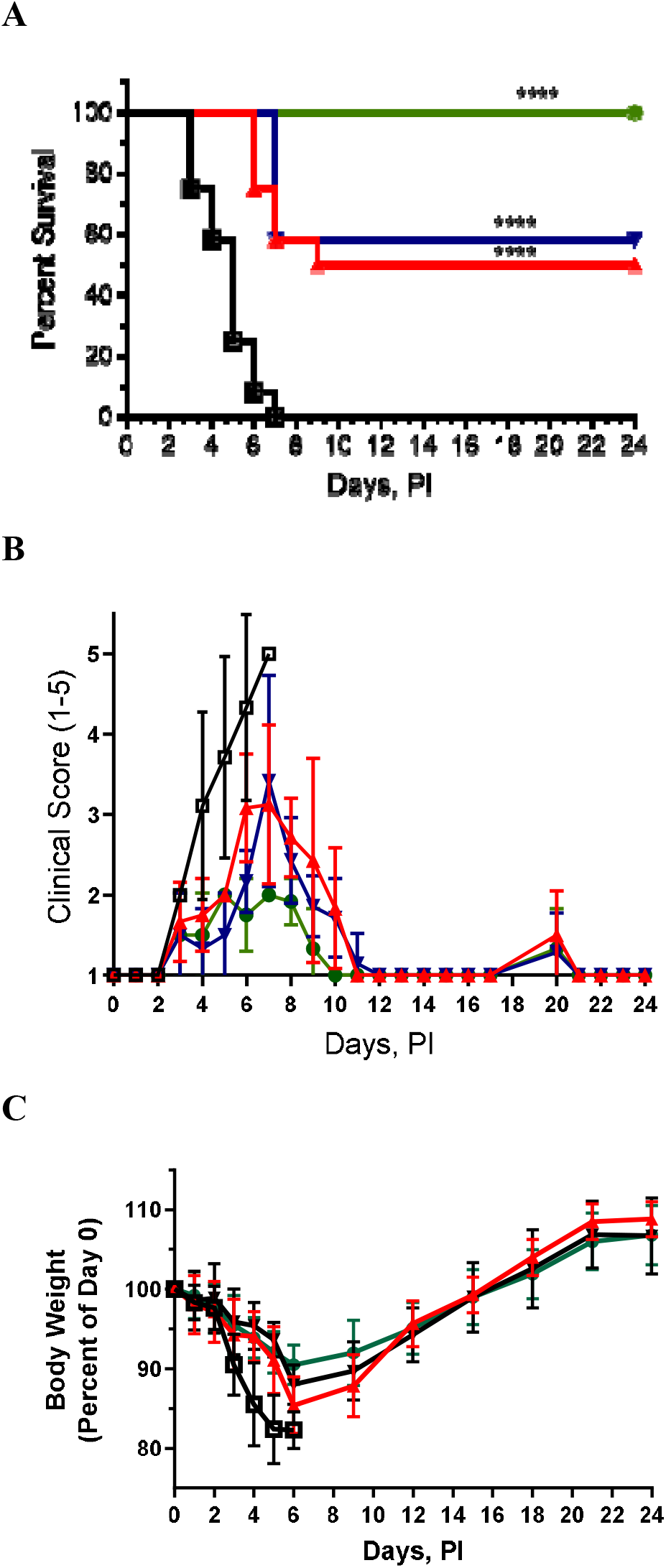
Effect of EIDD-1931 treatment initiated 2 hours before lethal Ebola virus challenge. (A) Kaplan-Meier survival curves; (B) Clinical score results; (C) Body weight dynamics. Vehicle-treated group (□); EIDD-1931 100 mg/kg QD treated group 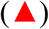; EIDD-1931 200 mg/kg QD group 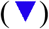; and EIDD-1931 200 mg/kg BID group 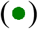. Data shown as average values ± standard deviation. **** p < 0.0001.

Vehicle-treated mice had a typical rapid increase in average clinical scores beginning on Study Day 3 and ending on Study Day 7 when all subjects had succumbed to EVD (**Fig. 2B**). The severity of EVD was significantly reduced in mice treated QD with 100 and 200 mg/kg EIDD-1931. The increase in average clinical score in mice from these two groups was similar between Study Days 3 through 7, and all surviving subjects no longer showed signs of EVD starting on Study Day 12. Mice treated with 200 mg/kg EIDD-1931 BID followed a similar score trend as mice receiving QD treatments but with lower maximum average values. Starting on Study Day 10, all mice in this group were noted as healthy.

Initial decreases in body weights were seen in all EBOV-infected groups (**Fig. 2C**). Vehicle-treated mice presented with a consistent decrease in average body weight beginning on Study Day 1. Once-daily EIDD-1931-treated mice also demonstrated decreases in average body weight beginning on Study Day 1 and continued through Study Day 6-9. Mice receiving 200 mg/kg BID also experienced initial body weight loss although the maximum decrease, which was observed on Study Day 6, was less than those in the vehicle and the QD treated groups. For all surviving EIDD-1931-treated mice, average body weights started to increase beginning on Study Days 6-9 and continued to increase through the end of the study (Study Day 24).

Blood samples for terminal serum viral loads from eight animals were available from mice receiving vehicle treatment (all euthanized in moribund conditions) and determined to have mean virus titers of 2×10^11^ GEq/mL (n=8) and infectious titers between 10^6^ and 10^7^ PFU/ml (n=5). Blood samples from all twelve mice receiving 200 mg/kg of EIDD-1931 BID (all survived to the end of the study) were available and determined to have viral titer loads below the lower limit of detection for the assay (LLOD) of 1×10^3^ GEq/mL. Three of 8 blood samples available from mice receiving EIDD-1931 at 100 mg/kg QD showed virus loads of 10^6^, 10^7^ and 10^10^ GEq/mL. It is interesting that mice with the 10^6^ and 10^7^ GEq/ml PCR titers (both survived to the end of the study) had no measurable infectious virus. The mouse with 10^10^ PCR titer, which succumbed on study day 6, had 10^6^ PFU/ml. The remaining five blood samples from this group had serum viral loads below the LLOD. One of these PCR-negative mice succumbed to EVD on study day 9 and the other four survived to the end of the study. Out of ten blood samples that were collected from mice receiving 200 mg/kg QD, seven were from mice survived to the end of the study and were determined to have serum viral loads below the LLOD. The three remaining samples were from mice that succumbed to disease on study day 7 and were determined to have viral loads of 10^8^ to 10^10^ GEq/mL and infectious titers of 10^4^ to 10^5^ PFU/ml.

### 3.2 Therapeutic treatment with EIDD-1931

Given the superior performance of BID dosing in prophylaxis studies, it was utilized in therapeutic studies. Mice in 3 treatment groups were orally administered vehicle or 200 mg/kg doses BID beginning 6 or 24 hours post virus challenge (**Table 1**, Study 2). Virus challenge dose in this study was determined to be higher than expected, ∼250 PFU/mouse (∼7500 LD_50_). All mice challenged with virus followed by BID dosing with vehicle succumbed to the disease by Study Days 2-6 (**Fig. 3A**). Eleven out of 12 (92%) mice treated with 200 mg/kg BID starting 6 hours post challenge survived to the end of the study (p<0.0001). Delay of treatment initiation by 24 hours post infection resulted in an increase of mean time-to-death by 1 day compared to the vehicle-treated group, although all mice succumbing to EVD. When 200 mg/kg BID dosing was initiated 48 hours post challenge, time-to-death and clinical course of EVD was no different than that observed for the vehicle-treated mice (data not shown).

**FIG 3.**
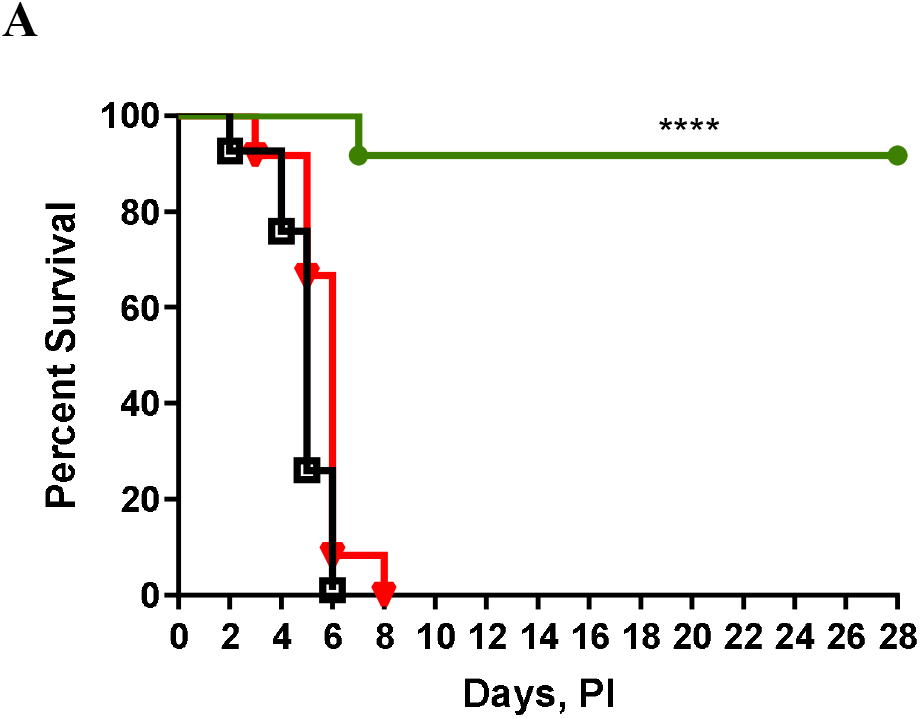
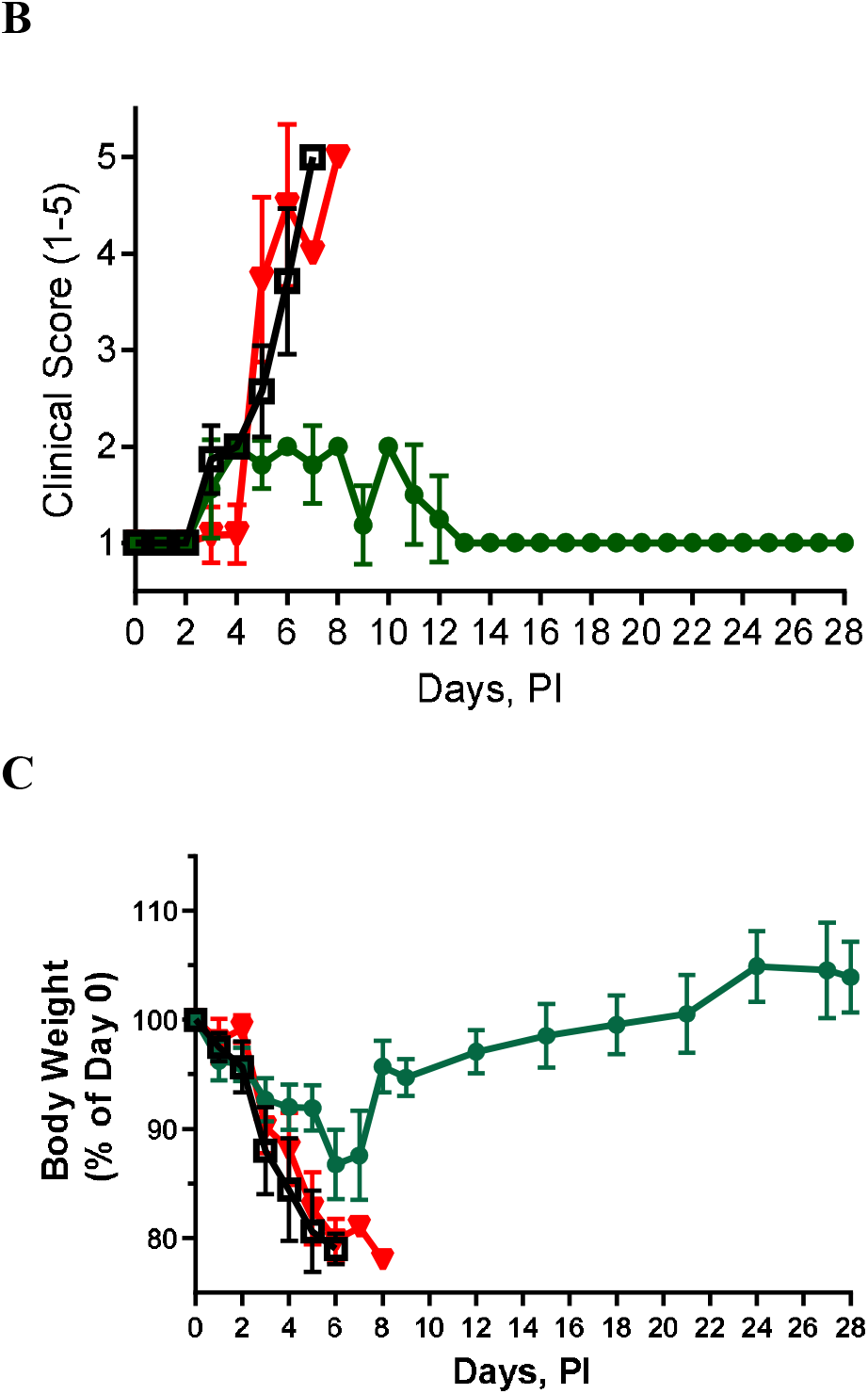
Effect of therapeutic EIDD-1931 treatment when initiated 6 or 24 hours post lethal Ebola virus-challenge. (A) Animal survival curves; (B) Clinical score results; (C) Body weight dynamics. Vehicle-treated mice (□), mice treated with EIDD-1931 200 mg/kg BID starting 6 hours post-infection 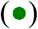, and mice treated with EIDD-1931 200 mg/kg BID starting 24 hours post-infection 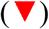. Data shown as average values ± standard deviation. **** p <0.0001.

Mice receiving treatment starting 6 hours post challenge had a minimal increase in average clinical score between Study Days 3-10 (**Fig. 3B**). The average clinical score decreased in this group after Day 7 and mice remained healthy thereafter. Vehicle and EIDD-1931 treated mice dosed starting 24 hours post challenge had rapid loss of body weight starting on Study Day 2-3, which continued until all animals in these groups succumbed to EVD **(Fig. 3C)**. Mice treated with EIDD-1931 starting 6 hours post challenge also showed an initial decrease in body weight that lasted through Study Day 7. Body weights began to recover after Day 7 in this group and returned to or exceeded the Study Day 0 values by the end of the study.

Results from qRT-PCR analysis of 6 terminal serum samples available from mice receiving vehicle treatment and euthanized on Study Days 2-6 indicated viral loads ranging from 8×10^9^ to 3×10^11^ GEq/mL and all had infectious virus load between 10^6^ and 10^9^ PFU/ml. Eleven analyzable terminal serum samples (all from surviving mice) were collected at the end of the study from mice receiving EIDD-1931 treatment initiated 6 hours post virus challenge. Nine of the samples had virus loads below the LLOD (1×10^3^ GEq/mL) while 2 other samples were above the LLOD but below the lower limit of quantification (LLOQ, 1×10^6^ GEq/mL). All these samples had undetectable infectious virus. Seven terminal serum samples were collected on Study Days 5-8 from mice receiving EIDD-1931 treatment initiated 24 hours post virus challenge. These samples had detectable and quantifiable levels of virus: one sample had 10^6^ GEq/mL and six other had 10^10^ to 10^11^ GEq/mL. All seven samples had detectable infectious virus, ranging from 10^2^ to 10^8^ PFU/ml.

## 4. DISCUSSION

Ebola virus infections result in high mortality and morbidity presenting a significant threat to human health. Expansion of human activities to areas of endemic Ebola disease coupled with the challenges of early detection and prompt responses makes future epidemics more likely to spread broadly and emerge in different, previously not affected, geographical locations. Despite the availability of monoclonal antibodies for the treatment of EBOV infections there is a clear need for an easily accessible, oral therapeutic that is broadly efficacious against multiple filoviruses and that overcomes the tissue penetration issues associated with monoclonal antibodies.

The availability of a robust small animal model of EVD that mimics disease in humans and can serve as a reliable first line system in which therapeutic agents and vaccines can be evaluated is critical. Mice are a preferred small animal for model research. Although the immunocompetent mouse model of EBOV infection is recognized for the failure to produce some key (though not constant) symptoms including coagulopathy, hemorrhagic signs, or petechial rash, this model successfully represents many of the hallmark characteristics of EVD in humans and NHPs including rapid onset of viremia, high viral burden and extensive necrosis in multiple organs, widespread apoptosis of lymphocytes, and a proinflammatory cytokine response (St Clair et al. 2017).

Anti-EBOV activity of NHC reported in cell culture models of infection (Reynard et al. 2015) supported the idea of conducting efficacy studies in the mouse model of lethal maEBOV infection. The pharmacokinetic and tissue distribution profile of EIDD-1931 had previously been determined in CD1 mice and showed that after oral administration EIDD-1931 was rapidly absorbed and widely distributed to tissues involved in EVD, including the spleen, liver, heart, lung, and brain (Painter et al., 2019). Since the bioavailability of EIDD-1931 in mice is high, there is no additional advantage of dosing with the prodrug molnupiravir (**Figure 1**); we have demonstrated that mouse plasma exposure to EIDD-1931 was very similar after oral dosing with EIDD-2801 or EIDD-1931 (Toots et al 2019). The distribution of EIDD-1931 to organs and the levels of tissue anabolism to its active 5’-triphosphate form are determined by circulating plasma levels of EIDD-1931 regardless of whether animals are dosed with molnupiravir or EIDD-1931.

The study described in this paper is the first to show that EIDD-1931 can effectively protect mice from death caused by EVD. Specifically, this data demonstrated that prophylactic oral dosing with EIDD-1931 resulted in a 100% survival rate (n=12, p<0.0001) when used in a BID dosing regimen. While average clinical scores were elevated and body weights decreased during initial stages of treatment, the clinical scores returned to normal and body weights increased steadily after days 7-9 post challenge. Oral treatment with EIDD-1931 initiated 6 and 24 hours post infection was also evaluated. In this experiment where mice were challenged with ∼7500 LD_50_ and started to die as early as on Day 2 post-challenge (**Fig. 3A**), the window for therapeutic initiation of treatment was found to be quite short: all mice died when treatment initiation was delayed by 24 or 48 hours post challenge. However, when EIDD-1931 treatment was initiated 6 hours post challenge, 92% of mice survived. Consistent with the prophylactic dosing results, average initial clinical scores were elevated, and body weights decreased during the initial phase of treatment, but the clinical scores subsequently returned to normal and body weights increased steadily until the end of the study. Protection of mice from a lethal challenge of maEBOV indicates that after oral dosing with EIDD-1931, adequate exposure to EIDD-1931-5’-triphosphate was achieved in tissues and organs critical to the pathogenesis of EVD. It should be noted that in human clinical trials, EIDD-1931 plasma exposure achieved after dosing with molnupiravir was well comparable to that achieved in mice after oral dosing with 200 mg/kg of EIDD-1931 (Painter et al. 2019, Painter et al. 2021, Khoo et al. 2021).

The assessment of molnupiravir in human patients as a potential prophylaxis and treatment for EVD should be facilitated by a well-established record of safety data from multiple clinical studies and from extensive use under Emergency Use Authorization (EUA) protocols for the treatment of COVID-19. (Painter et al. 2021, Arribas et al. 2021, Bernal et al. 2021, FDA News Release 12/23/2021).

The result of the PALM study (Mulangu et al 2019) has been a major advance in the Ebola therapy using mAbs, however the residual mortality of 34% despite treatment suggests there is room for improvement, including combination treatment with antivirals. The direct-acting antiviral molnupiravir could be studied as an add-on therapy to standard-of-care to evaluate for reduction in mortality and viral shedding. Another potential indication for molnupiravir could include post-exposure prophylaxis for highly exposed Ebola contacts, particularly those located in remote areas where facilities for intravenous infusion of mAbs are limited. Even when such capabilities are available, the community contacts are often reluctant to go the facility for IV infusion because of stigma and also because they don’t feel that they are sick. Oral pills taken at home could be an attractive alternative. The in vitro and in vivo activity against EBOV coupled with good bioavailability of molnupiravir supports further testing and clinical development using an NHP model of Ebola infection, a model the US FDA agrees is relevant and adequately characterized to support filing under the Animal Efficacy Rule (FDA 21CFR314). Preliminary PK and tissue distribution studies conducted in Cynomolgus macaques have shown that EIDD-1931 is widely distributed (data not shown), including into brain and testes, that may sequester virus and are untreated by antibody therapies.

## ABBREVIATIONS

EBOV: Ebola virus
EVD: Ebola virus disease
NHC: *N*^*4*^-Hydroxycytidine
PFU: plaque forming units
LD_50,_: virus dose causing 50% lethality
maEBOV: mouse-adapted Ebola virus
QD: once daily
BID: twice daily
PO: oral dosing
IV: intravenous infusion
LLOD: low limit of detection
LLOQ: low limit of quantitation
GEq: genome equivalent
mAb: monoclonal antibody

## Acknowledgements

Emory University has utilized the Preclinical Services offered by the National Institute of Allergy and Infectious Diseases (NIAID), National Institutes of Health (NIH), Department of Health and Human Services.

## REFERENCES

Agostini ML, Pruijssers AJ, Chappell JD, Gribble J, Lu X, Andres EL, Bluemling GR, Lockwood MA, Sheahan TP, Sims AC, Natchus MG, Saindane M, Kolykhalov AA, Painter GR, Baric RS, Denison MR. 2019. Small molecule antiviral β-D-N4-hydroxycytidine inhibits a proofreading-intact coronavirus with a high genetic barrier to resistance. J Virol 93(24), e01348–19. https://doi.org/10.1128/JVI.01348-19

Arribas et al. 2021. Randomized Trial of Molnupiravir or Placebo in Patients Hospitalized with Covid-19. NEJM Evidence. DOI: https://evidence.nejm.org/doi/10.1056/EVIDoa2100044

Bausch DG. 2021. The need for a new strategy for Ebola vaccination. Nat Med 27(4), 580–581. https://doi.org/10.1038/s41591-021-01313-w

Jayk Bernal A, Gomes da Silva MM, Musungaie DB, Kovalchuk E, Gonzalez A, Delos Reyes V, Martín-Quirós A, Caraco Y, Williams-Diaz A, Brown ML, D. J, Pedley A, Assaid C, Strizki J, Grobler JA, Shamsuddin HH, Tipping R, Wan H, Paschke A, Butterton JR, Johnson MG, De Anda C; MOVe-OUT Study Group. 2021. Molnupiravir for oral treatment of Covid-19 in nonhospitalized patients. N Engl J Med, NEJMoa2116044. https://doi.org:10.1056/NEJMoa2116044

Bray M, Davis K, Geisbert T, Schmaljohn C, Huggins J. 1998. A mouse model for evaluation of prophylaxis and therapy of Ebola hemorrhagic fever. J Infect Dis 178, 651–661. https://doi.org/10.1086/515386

Bwaka MA, Bonnet M-J, Calain P, Colebunders R, De Roo A, Guimard Y, Katwiki KR, Kibadi K, Kipasa MA, Kuvula KJ, Mapanda BB, Massamba M, Mupapa KD, Muyembe-Tamfum J-J, Ndaberey E, Peters CJ, Rollin PE, Van den Enden E. 1999. Ebola hemorrhagic fever in Kikwit, Democratic Republic of the Congo: clinical observations in 103 patients. J Infect Dis 179, S1–S7. https://doi.org/10.1086/514308

Den Boon S, Marston B J, Nyenswah T G, Jambai A, Barry M, Keita S, Dye C. 2019. Ebola virus infection associated with transmission from survivors. Emerg Infect Dis 25(2), 240–246. https://doi.org/10.3201/eid2502.181011

EMA Web Page. 2019. First vaccine to protect against Ebola. 18 Oct 2019. Accessed on 08 Oct 2019. Available from: https://www.ema.europa.eu/en/news/first-vaccine-protect-against-ebola

FDA (US Food and Drug Administration) Code of Federal Regulations (CFR) Title 21, Sec 314. Applications for FDA Approval to market a new drug, subpart i, approval of new drugs when human efficacy studies are not ethical or feasible. Accessed 7 Feb 2022. Available from: https://www.accessdata.fda.gov/scripts/cdrh/cfdocs/cfcfr/CFRSearch.cfm?CFRPart=314&showFR=1&subpartNode=21:5.0.1.1.4.9

FDA (US Food and Drug Administration) News Release Web Page.2019. First FDA-approved vaccine for the prevention of Ebola virus disease, marking a critical milestone in public health preparedness and response. 19 Dec 2019. Accessed 08 Oct 2021. Available from: https://www.fda.gov/news-events/press-announcements/first-fda-approved-vaccine-prevention-ebola-virus-disease-marking-critical-milestone-public-health

FDA (US Food and Drug Administration) News Release, December 23, 2021. Coronavirus (COVID-19) Update: FDA Authorizes Additional Oral Antiviral for Treatment of COVID-19 in Certain Adults. https://www.fda.gov/news-events/press-announcements/coronavirus-covid-19-update-fda-authorizes-additional-oral-antiviral-treatment-covid-19-certain

Feldmann H, Geisbert TW. 2011. Ebola haemorrhagic fever. Lancet 377, 849–862. https://doi.org/10.1016/S0140-6736(10)60667-8

Fischer WA 2nd, Eron JJ Jr, Holman W, Cohen MS, Fang L, Szewczyk LJ, Sheahan TP, Baric R, Mollan KR, Wolfe CR, Duke ER, Azizad MM, Borroto-Esoda K, Wohl DA, Coombs RW, James Loftis A, Alabanza P, Lipansky F, Painter WP. 2022. A phase 2a clinical trial of molnupiravir in patients with COVID-19 shows accelerated SARS-CoV-2 RNA clearance and elimination of infectious virus. Sci Transl Med 14(628), eabl7430. https://doi.org/10.1126/scitranslmed.abl7430. Epub 2022 Jan 19.

Gordon CJ, Tchesnokov EP, Schinazi RF, Götte M. 2021. Molnupiravir promotes SARS-CoV-2 mutagenesis via the RNA template. J Biol Chem 297(1),100770. https://doi.org/10.1016/j.jbc.2021.100770

Haddock E, Feldmann H, Marzi A. 2018. Ebola virus infection in commonly used laboratory mouse strains. J Infect Dis 218(Suppl_5), S453–S457. https://doi.org/10.1093/infdis/jiy208

Keita AK, Koundouno FR, Faye M, Düx A, Hinzmann J, Diallo H, Ayouba A, Le Marcis F, Soropogui B, Ifono K, Diagne MM, Sow MS, Bore JA, Calvignac-Spencer S, Vidal N, Camara J, Keita MB, Renevey A, Diallo A, Soumah AK, Millimono SL, Mari-Saez A, Diop M, Doré A, Soumah FY, Kourouma K, Vielle NJ, Loucoubar C, Camara I, Kourouma K, Annibaldis G, Bah A, Thielebein A, Pahlmann M, Pullan ST, Carroll MW, Quick J, Formenty P, Legand A, Pietro K, Wiley MR, Tordo N, Peyrefitte C, McCrone JT, Rambaut A, Sidibé Y, Barry MD, Kourouma M, Saouromou CD, Condé M, Baldé M, Povogui M, Keita S, Diakite M, Bah MS, Sidibe A, Diakite D, Sako FB, Traore FA, Ki-Zerbo GA, Lemey P, Günther S, Kafetzopoulou LE, Sall AA, Delaporte E, Duraffour S, Faye O, Leendertz FH, Peeters M, Toure A, Magassouba NF. 2021. Resurgence of Ebola virus in 2021 in Guinea suggests a new paradigm for outbreaks. Nature 597, 539–543. https://doi.org/10.1038/s41586-021-03901-9

Kortepeter MG, Bausch DG, Bray M. 2011. Basic clinical and laboratory features of filoviral hemorrhagic fever. J Infect Dis 204, S810–S816. https://doi.org/10.1093/infdis/jir299

Mulangu S, Dodd LE, Davey RT Jr, Tshiani Mbaya O, Proschan M, Mukadi D, Lusakibanza Manzo M, Nzolo D, Tshomba Oloma A, Ibanda A, Ali R, Coulibaly S, Levine AC, Grais R, Diaz J, Lane HC, Muyembe-Tamfum JJ, PALM Writing Group, Sivahera B, Camara M, PALM Consortium Study Team. 2019. A randomized, controlled trial of Ebola virus disease therapeutics. NEJM, 381(24), 2293–2303. https://doi.org/10.1056/NEJMoa1910993

Painter GR, Bowen RA, Bluemling GR, DeBergh J, Edpuganti V, Gruddanti PR, Guthrie DB, Hager M, Kuiper DL, Lockwood MA, Mitchell DG, Natchus MG, Sticher ZM, Kolykhalov AA. 2019. The prophylactic and therapeutic activity of a broadly active ribonucleoside analog in a murine model of intranasal Venezuelan equine encephalitis virus infection. Antiviral Res 171, 104597. https://doi.org/10.1016/j.antiviral.2019.104597

Painter WP, Holman W, Bush JA, Almazedi F, Malik H, Eraut Ncje, Morin MJ, Szewczyk LJ, Painter GR. 2021. Human safety, tolerability, and pharmacokinetics of molnupiravir, a novel broad-spectrum oral antiviral agent with activity against SARS-CoV-2. Antimicrob Agents Chemother 65(5), e02428–20. https://doi.org/10.1128/AAC.02428-20

Rewar S, Mirdha D. 2014. Transmission of ebola virus disease: an overview. Ann Glob Health, 80(6), 444–451. https://doi.org/10.1016/j.aogh.2015.02.005

Reynard O, Nguyen X-N, Alazard-Dany N, Barateau V, Cimarelli A, Volchkov V. 2015. Identification of a New Ribonucleoside Inhibitor of Ebola Virus Replication. Viruses 7:2934.33. https://doi.org/10.3390/v7122934

Sheahan TP, Sims AC, Zhou S, Graham RL, Pruijssers AJ, Agostini ML, Leist SR, Schäfer A, Dinnon KH, Stevens LJ, Chappell JD, Lu X, Hughes TM, George AS, Hill CS, Montgomery SA, Brown AJ, Bluemling GR, Natchus MG, Saindane M, Kolykhalov AA, Painter G, Harcourt J, Tamin A, Thornburg NJ, Swanstrom R, Denison MR, Baric RS. 2020. An orally bioavailable broad-spectrum antiviral inhibits SARS-CoV-2 in human airway epithelial cell cultures and multiple coronaviruses in mice. Sci Transl Med 12, eabb5883. https://doi.org/10.1126/scitranslmed.abb5883

Shurtleff AC, Bloomfield HA, Mort S, Orr SA, Audet B, Whitaker T, Richards MJ, Bavari S. 2016. Validation of the filovirus plaque assay for use in preclinical studies. Viruses 8(4), 113. https://doi.org/10.3390/v8040113

St Claire MC, Ragland DR, Bollinger L, Jahrling PB. 2017. Animal models of Ebolavirus infection. Comp Med 67, 253–262. PMID: 28662754; PMCID: PMC5482517

Taylor K. 2007. Biodefense: research methodology and animal models. Emerg Infect Dis 13(3), 523. https://doi.org/10.3201/eid1303.061488

Toots M, Yoon J-J, Cox RM, Hart M, Sticher ZM, Makhsous N, Plesker R, Barrena AH, Reddy PG, Mitchell DG, Shean RC, Bluemling GR, Kolykhalov AA, Greninger AL, Natchus MG, Painter GR, Plemper RK. 2019. Characterization of orally efficacious influenza drug with high resistance barrier in ferrets and human airway epithelia. Sci Transl Med 11, eaax5866. https://doi.org/10.1126/scitranslmed.aax5866

Urakova N, Kuznetsova V, Sokratian A, Frolova EI, Frolov I, Crossman DK, Crowley MR, Guthrie DB, Kolykhalov AA, Lockwood MA, Natchus MG, Painter GR, Painter GR. 2018. β-d-N(4)-Hydroxycytidine Is a potent anti-alphavirus compound that induces a high level of mutations in the viral genome. J Virol 92, e01965–17. https://doi.org/10.1128/JVI.01965-17

Vetter P, Fischer Iiwa, Schibler M, Jacobs M, Bausch DG, Kaiser L. 2016a. Ebola virus shedding and transmission: review of current evidence. J Infect Dis 214, S177-S184. doi.org/10.1093/infdis/jiw254

Vetter P, Kaiser L, Schibler M, Ciglenecki I, Bausch DG. 2016b. Sequelae of Ebola virus disease: the emergency within the emergency. Lancet Infect Dis 16, e82–e91. https://doi.org/10.1016/S1473-3099(16)00077-3

WHO (World Health Organization) Disease Outbreak News. 2020. Ebola – African Region (AFRO), Democratic Republic of the Congo. 26 June, 2020. Accessed on 7 Feb 2022. Available from: https://www.who.int/emergencies/disease-outbreak-news/item/2020-DON284

Yoon J-J, Toots M, Lee S, Lee M-E, Ludeke B, Luczo JM, Ganti K, Cox RM, Sticher ZM, Edpuganti V, Mitchell DG, Lockwood MA, Kolykhalov AA, Greninger AL, Moore ML, Painter GR, Lowen AC, Tompkins SM, Fearns R, Natchus MG, Plemper RK. 2018. Orally efficacious broad-spectrum ribonucleoside analog inhibitor of influenza and respiratory syncytial viruses. Antimicrob Agents Chemother 62:e00766–18. https://doi.org/10.1128/AAC.00766-18

